# Cell-free prototyping strategies for enhancing the sustainable production of polyhydroxyalkanoates bioplastics

**DOI:** 10.1101/225144

**Authors:** Richard Kelwick, Luca Ricci, Soo Mei Chee, David Bell, Alexander J. Webb, Paul S. Freemont

**Author notes:** Joint First Authors (Richard Kelwick, Luca Ricci). To whom correspondence should be addressed: Paul Freemont, Section of Structural Biology Department of Medicine, Sir Alexander Fleming Building, South Kensington Campus, Exhibition Road London, SW7 2AZ UK.

## Abstract

The polyhydroxyalkanoates are a group of microbially-produced biopolymers that have been proposed as sustainable alternatives to several oil-derived plastics. However, polyhydroxyalkanoates are currently more expensive to produce than oil-derived plastics and therefore, more efficient production processes would be desirable. Cell-free transcription-translation-based metabolic engineering strategies have been previously used to optimise several different biosynthetic pathways but not the polyhydroxyalkanoates biosynthetic pathways. Here we have developed several *Escherichia coli* cell-free transcription-translation-based systems for *in vitro* prototyping of polyhydroxyalkanoates biosynthetic operons, and also for screening relevant metabolite recycling enzymes. These cell-free transcription-translation reactions were customised through the addition of whey permeate, an industrial waste that has been previously used as a low-cost feedstock for optimising *in vivo* polyhydroxyalkanoates production. We found that the inclusion of an optimal concentration of whey permeate enhanced relative cell-free GFPmut3b production by ~20% compared to control reactions that did not include whey permeate. An analysis of pH in our cell-free reactions suggests that the observed increase in GFPmut3b production was likely through enhanced ATP generation, as a consequence of the glycolytic processing of lactose present in whey permeate. We also found that whey permeate enhanced cell-free reactions produced ~3μM (R)-3HB-CoA, whilst, coupled cell-free biotransformation/transcription-translation reactions produced a ten-fold greater yield of (R)-3HB-CoA. These reactions were also used to characterise a *Clostridium propionicum* propionyl CoA transferase enzyme that can recycle Acetyl-CoA. Together our data demonstrate that cell-free approaches can be used to complement *in vivo* workflows for identifying additional strategies for optimising polyhydroxyalkanoates production.

## 1. Introduction

The mass production of oil-derived plastics has resulted in widespread and potentially irreversible global ecological damage (Eriksen et al., 2014; Suaria et al., 2016). Despite these environmental consequences, oil-derived plastics are still in very high-demand, due to their versatility and low-cost of production, that even with improvements in recycling processes, *de novo* plastic production has not been reduced (Geyer et al., 2017). Moreover, the societal and technological challenges that are associated with the introduction of more sustainable alternatives to oil-derived plastics are significant, several of which have been extensively reviewed (Kishna et al., 2017; Kourmentza et al., 2017). Synthetic biology is a field that is driven by the development and carefully-considered implementation of rationally-engineered biotechnologies that might help to address local or global challenges (Kelwick et al., 2015a, 2014; Tait, 2017; Webb et al., 2017, 2016). Developments in synthetic biology, along with a continuum of advancements in metabolic engineering, could potentially enable the routine and highly scalable production of microbially-produced biopolymers, such as polyhydroxyalkanoates (PHAs) (Chen et al., 2017; Gustavsson and Lee, 2016; Kelwick et al., 2015b; Lee, 2006; Meng and Chen, 2017). PHAs share several material characteristics with some of the most widely used oil-derived plastics, but beneficially, PHAs are also biodegradable (Dietrich et al., 2017; Z. Li et al., 2016). In order to develop PHAs as viable alternatives to oil-derived plastics, great efforts have been undertaken to design more efficient microbial PHAs production processes through the rational engineering of biosynthetic pathways (Hiroe et al., 2012; Kelwick et al., 2015b; T. Li et al., 2016; Tao et al., 2017), metabolite recycling processes (e.g. Acetyl-CoA) (Beckers et al., 2016; Ken’Ichiro Matsumoto et al., 2013) and the use of industrially-sourced, low-cost feedstocks (e.g. whey permeate) (Ahn et al., 2000; Cui et al., 2016; Kim, 2000; Nielsen et al., 2017; Nikel et al., 2006; Wong and Lee, 1998). Alternative, enzymatic approaches for the production of PHAs have also been investigated (Han et al., 2011; Opgenorth et al., 2016; Thomson et al., 2010).

From these extensive studies, we reasoned that cell-free transcription-translation (TX-TL) systems could be used to efficiently characterise different PHA biosynthetic operons. Indeed, cell-free TX-TL systems have been widely used as prototyping platforms for characterising DNA based parts, devices and systems including: DNA regulatory elements (Chappell et al., 2013; Garamella et al., 2016; Kelwick et al., 2016), increasingly complex genetic circuits (Hori et al., 2017; Niederholtmeyer et al., 2015; Shin and Noireaux, 2012; Sun et al., 2014; Takahashi et al., 2015) and cell-free medical bioreporter devices (Pardee et al., 2014; Slomovic et al., 2015; Wen et al., 2017). These cell-free applications have also renewed interest in cell-free metabolic engineering approaches for *in vitro* enzyme screening and prototyping of entire biosynthetic pathways (Dudley et al., 2015; Goerke et al., 2008; Hodgman and Jewett, 2012; Jewett et al., 2008; Karim and Jewett, 2016; Pardee et al., 2016; Ullah et al., 2016). Importantly, extract-based cell-free systems enable the *in situ* characterisation of enzymes and biosynthetic pathways within a metabolic context that is representative of the production strain (Karim et al., 2016). Since cell extracts contain not only the TX-TL machinery needed to express a biosynthetic operon but also relevant enzyme co-factors and potentially competing metabolic pathways, cell-free metabolic engineering approaches can be used to identify novel strategies that improve the *in vivo* activities of biosynthetic pathways (Perez et al., 2016). Yet, interestingly cell-free TX-TL systems have not been previously used to characterise PHA biosynthetic pathways. Here we report the development of several cell-free approaches for prototyping PHA biosynthetic operons that could be used in combination with *in vivo* strategies for optimising PHAs production.

## 2. Materials and Methods

### 2.1 Bacterial strains, constructs and growth conditions

All constructs and bacterial strains used in this study are listed in Supplementary Table 1. *E. coli* NEB10-beta was used for cloning, whilst *E. coli* MG1655 was used for experiments. For plasmid recovery *E. coli strains* were grown in Luria-Bertani (LB) supplemented with either 34 μg/ml Chloramphenicol (final concentration), 100 μg/ml Ampicillin (final concentration) or 50 μg/ml Kanamycin (final concentration), as appropriate, at 37°C, with shaking (220 rpm).

### 2.2 Construct assembly

The empty vector negative control constructs EV_101 (BBa_J23101; pRK21) and EV_104 (BBa_K608002; pRK4) were originally sourced from the 2013 distribution of the iGEM Registry of Standard Biological Parts (partsregistry.org). EV_104 was transformed into chemically competent *E. coli* MG1655 to create strain S_RK001. *phaCAB* bioplastic operon constructs: The native-phaCAB construct (BBa_K934001; pRK51) was transformed into *E. coli* MG1655 to create strain, S_RK002. The constitutive 104 *phaCAB* operon construct (C104 - BBa_K1149052; pRK61) was transformed into *E. coli* MG1655 to create strain, S_RK003. The native (pRK51) and C104 (pRK61) *phaCAB* constructs are also available in the iGEM Registry of Standard Biological Parts (partsregistry.org) and were also reported in our previous study (Kelwick et al., 2015b). To generate a constitutive phaCAB-operon with an inactive PhaC enzyme, PCR reactions were undertaken using PCR primer pairs PhaCCys319Ala_F (RK007) and PhaCCys319Ala_R (RK008) with C104 (pRK61) as the DNA template. The resultant PCR product was digested with DpnI restriction enzyme, phosphorylated, self-ligated (Quick Ligase - New England Biolabs, UK) and transformed into *E. coli* NEB10-beta, resulting in *phaCAB*-operon strain/plasmid, C104ΔPhaC_C319A (pRK7). This plasmid was then purified from *E. coli* NEB10-beta using the Qiaprep Spin Miniprep Kit (Qiagen, Germany) and transformed into *E. coli* MG1655 competent cells, resulting in *E. coli* MG1655 strain S_RK004.

To generate the GFP expression plasmid (101_GFP), plasmid pRK1 was digested with XbaI and PstI to generate a fragment containing the RBS B0034, *gfpmut3b* and terminators B0010 and B0012. This fragment was ligated (Quick Ligase - New England Biolabs, UK) into a SpeI and PstI digested EV_101 backbone (pRK3) and transformed into *E. coli* NEB10-beta to create plasmid/strain, pLR036. pT7 constructs: A negative control, empty vector, encoding a T7 promoter (BBa_I719005; pRK9) was originally sourced from the 2013 distribution of the iGEM Registry of Standard Biological Parts (partsregistry.org).

Plasmid pT7_PCT (pRK12) was generated to enable inducible expression of the propionyl CoA transferase (PCT) gene from *Clostridium propionicum*. To generate the construct the *pct* gene sequence was obtained from GenBank (AJ276553.1) and then synthesised as a GeneArt String DNA fragment (Life Technologies, USA). The GeneArt String DNA fragment was blunt-end cloned, using the Zero Blunt PCR Cloning Kit (Life Technologies, USA), into pCR-Blunt vector according to the manufacturer’s instructions – producing plasmid (pRK11). Afterwards, a PCR reaction was carried out using primers F1_PCT-FW (RK011) and F1_PCT-RV (RK012), where plasmid pRK11 served as the DNA template. The resultant PCR product generated a *pct* encoding DNA fragment. Additionally, a PCR reaction was carried out using primers T7_INF_FWD (RK009) and T7_INF_REV (RK010), where plasmid pRK10 served as the DNA template. The resultant PCR product generated a linearised plasmid vector. The PCR products from both reactions were used in an In-Fusion cloning reaction (Takara Bio, USA) and 2.5 μl of the completed reaction was transformed into NEB10-beta producing strain/plasmid pRK12 The DNA sequences of all inserts/constructs were verified by the sequencing service provided by Eurofins Genomics GmbH (Ebersberg, Germany). Primers used for sequencing and cloning are listed in Supplementary Table 2.

### 2.3 Preparation of cell extracts

To prepare cell-free extracts, *E. coli* MG1655 cells were revived from glycerol stocks onto 2x YTP plates (31 g/L 2x YT, 40 mM potassium phosphate dibasic, 22 mM potassium phosphate monobasic and 15 g/L agar). Once streaked, plates were incubated overnight at 37°C. Individual colonies were inoculated into 5 ml 2x YTP medium (31 g/L 2x YT, 40 mM potassium phosphate dibasic and 22 mM potassium phosphate monobasic) and incubated overnight with shaking (220 rpm) at 37°C. 2.5 ml aliquots of the resultant cultures were used to inoculate flasks containing 50 ml 2x YTP medium. These cultures were then incubated at 37°C with shaking (220 rpm) until the cell density reached an 0D600_nm_ of 2. Finally, 25 ml of the resultant cultures were used to inoculate flasks containing 500 ml 2x YTP medium. These cultures were subsequently incubated at 37°C with shaking (220 rpm) until cell density reached an 0D600_nm_ of between 2 to 3. For the preparation of cell extracts from strain S_RK004, the cells were cultured using these same conditions except that 2x YTP plates and media were also supplemented with 34 μg/ml Chloramphenicol.

To harvest cells, 500 ml cultures were centrifuged at 3,220 *g* for 15 minutes. Cell pellets were re-suspended into 20 ml S30-A buffer (14 mM Magnesium (Mg) glutamate, 60 mM Potassium (K) glutamate, 50 mM Tris, 2mM DTT, pH 7.7) and transferred into a pre-weighed 50 ml Falcon tube. Each 50 ml Falcon tube was centrifuged (2,000 g, 10 min, 4°C), pellets washed with 20 ml S30-A buffer and subsequently re-centrifuged (2,000 g, 10 min, 4°C) to form the final cell pellets in preparation for cell lysis. To determine the weight of the cell pellet, the weight of the 50 ml falcon tube was subtracted from the combined weight of the 50 ml tube and cell pellet. Pellets were stored at -80°C for no more than 48 h, prior to cell lysis.

To lyse the cells, pellets were defrosted on ice and re-suspended into 1 ml S30-A buffer per gram of cell pellet and aliquoted as 1 ml samples in 1.5 ml microtubes. Samples were sonicated on ice (3 x 40 seconds with 1-minute cooling interval; output frequency: 20 KHz; amplitude: 50%) and then centrifuged (12,000 *g* at 4°C for 10 min). The supernatants were removed, aliquoted at 500 μl into 2 ml screw cap tubes and incubated with shaking (220 rpm) at 37°C for 80 min. Post pre-incubation, samples were stored on ice and then centrifuged (12,000 *g* at 4 °C for 10 min). Supernatants were removed and were aliquoted into dialysis cassettes (GeBAflex-Maxi Dialysis Tubes - 8 kDa MWCO, Generon) for dialysis into S30-B buffer (14 mM Mg-glutamate, 60 mM K-glutamate, ~5 mM Tris, 1mM DTT; pH 8.2) with stirring at 4°C for 3 hr. Post-dialysis samples were centrifuged (12,000 *g* at 4 °C for 10 min), the extract supernatants were aliquoted into 1.5 ml tubes, flash frozen in liquid nitrogen and stored at -80°C for use in cell-free reactions.

### 2.4 Cell-free transcription-translation reactions

Cell-free transcription-translation reactions consisted of three parts mixed together in the indicated ratios: cell extract (33% v/v), energy buffer (42% v/v) and plasmid DNA (25% v/v). The final reaction conditions were: 8 mM Mg-glutamate, 260 mM K-glutamate, 1.5 mM each amino acid (except leucine - 1.25 mM leucine), 1.5 mM of both ATP and GTP, 0.9 mM of both CTP and UTP, 1.5 mM spermidine, a range (0-5.748 g/L) of Molkolac instant demineralised whey permeate (Orchard Valley Food Ingredients, UK), 0 or 100 μM (final concentration) SNARF-5F pH sensitive dye (Invitrogen, USA) and 10 nM (final concentration) plasmid DNA. For analysis of cell-free GFP production or pH, 10 μl cell-free reactions were aliquoted into individual wells of 384-well plates (Griener bio-one, NC, USA) and measured using a Clariostar plate reader (BMG, UK) with the following settings: GFP - excitation 483-14 nm and emission 530-30 nm, for pH - excitation 514 nm and the ratio of two different emissions, 580/640 nm were measured and used in conjunction with a calibration curve (Supplementary Figure 1). Plates were sealed, shaken prior to each reading cycle (500 rpm) and the plate reader was set to incubate the cell-free reactions at 30°C. For gas chromatography-mass spectrometry (GC-MS) analysis of 3HB content in cell-free transcription-translation reactions, the reactions were scaled up to 30 μl and were incubated in 1.5 ml tubes at 30°C for 5 h. Cell-free transcription-translation reactions were then treated with 30 μl ice cold acetonitrile, centrifuged (12,000 *g* at 4 °C for 10 min) and the resultant supernatants were used for downstream GC-MS analysis. For analysis of Acetyl-CoA content in cell-free transcription-translation reactions the reactions were scaled up to 30 μl and were incubated in 1.5 ml tubes at 30°C for 1 h. These samples were then deproteinised according to the manufacturer’s instructions in the PicoProbe Acetyl CoA Fluorometric Assay Kit (Abcam, UK). Briefly, cell-free samples were deproteinised using perchloric acid then neutralised with 3 M KHCO_3_ as per the kits instructions.

### 2.5 Cell-free biotransformation reactions

For GC-MS analysis of 3HB content in cell-free biotransformation reactions, 30 μl cell-free reactions were setup and consisted of three parts mixed together in the indicated ratios: cell extract (33% v/v), energy buffer (42% v/v) and plasmid DNA (25% v/v). The final reaction conditions were: 8 mM Mg-glutamate, 260 mM K-glutamate, 1.5 mM each amino acid (except leucine - 1.25 mM leucine), 1.5 mM of both ATP and GTP, 0.9 mM of both CTP and UTP, 1.5 mM spermidine, 0 or 0.004 g/L Molkolac instant demineralised whey permeate (Orchard Valley Food Ingredients, UK), 0 or 25 units T7 RNA polymerase and 0 or 10 nM (final concentration) of pT7_PCT (pRK12) plasmid DNA. Cell-free biotransformation reactions were incubated in 1.5 ml Eppendorf tubes at 30°C for 0 or 5 h at 30°C. The cell-free biotransformation reactions were then treated with 30 μl ice cold acetonitrile, centrifuged (12,000 *g* at 4 °C for 10 min) and the resultant supernatants were used for downstream GC-MS analysis. For analysis of acetyl-CoA content in cell-free biotransformation reactions, 30 μl reactions were incubated in 1.5 ml tubes at 30°C for 0 or 2.5h. These samples were then deproteinised according to the manufacturer’s instructions in the PicoProbe Acetyl CoA Fluorometric Assay Kit (Abcam, UK). Briefly, cell-free samples were deproteinised using perchloric acid then neutralised with 3 M KHCO3 as per the kits instructions.

### 2.6 SNARF-5F pH calibration curve

A pH-sensitive calibration curve was generated using the SNARF-5F dye, 5-(and-6)-carboxylic acid (Invitrogen, USA). The dye was dissolved in DMSO at 1 mM and stored at +4° C. SNARF-5F was diluted to 100 μM (final concentration) in 10 μl (total volume) of a range of different 100 mM Tris buffers (final concentration) that were set, using 0.5 M acetic acid, at a range of defined pH strengths. 10 μl aliquots of these mixtures were aliquoted into individual wells of a 384-well plate (Greiner bio-one, NC, USA). End-point fluorescence measurements were carried out using a Clariostar plate reader (BMG, UK) set to an excitation of 514 nm and a ratio of two different emissions, 580/640 nm. From the calibration curve a third order polynomial fitting was developed to extrapolate the pH values from the fluorescence 580/640 nm emissions ratio. These calibration curve data are shown in Supplementary Figure 1.

### 2.7 3-hydroxybutyric acid detection in cell-free samples using GC-MS

The cell free samples were subjected to trimethylsilylation reaction for the detection of 3-hydroxybutyric acid (3-HB) production using GC-MS. The 3-HB samples were centrifuged for 5 mins at 13,500 rpm, then 40 μl of the supernatant was dried under a gentle nitrogen stream. To the dried samples, 90 μl of MSTFA was added + MSTFA 1% TMCS silylation reagent (Thermo Fisher, part number: 11567851) and left to react at 37 °C for 30 minutes.

The commercial 3-HB standard (Sigma Aldrich, USA #54965-10G-F) was dissolved in methanol (Sigma Aldrich) at several different concentrations between the range of 0 μM to 200 μM in serial dilutions to generate a calibration curve (Supplementary Figure 2). 10 μl of the standards were then evaporated to dryness under a gentle nitrogen stream. The dried standards were treated with the trimethylsilylation reaction as described above.

The trimethylsilylated derivatised samples were analysed by GC-MS, using an Agilent Technologies 7890B GC and MSD 5977 series system with electron ionisation in SIM mode by monitoring ion with m/z value 117.1, 147.1, 191.1 and 233.0. Helium was used as the carrier gas. The temperatures of the injector and MS transfer line were 240°C and 250°C respectively, whereas MS quadrupole and MS source were 150°C and 250°C respectively. The samples were analysed with an injection volume of 1 μL, at a split ratio of 10 to 1. A temperature program was used for separation of the trimethylsilylated 3HB: initial temperature is 80°C, temperature increase 30°C/minute until 123°C, temperature increase 1°C/minute until 128°C, temperature increase 60°C/minute until 280°C, and hold for 3 minutes, followed by 0.5 minutes post run at 80°C.

### 2.8 Acetyl-CoA Assay

Acetyl-CoA content was quantified using the PicoProbe acetyl-CoA assay kit (ab87546, Abcam, UK). The 0-100 pM range acetyl-CoA standard curve was generated with a correlation coefficient of 0.9982 (Supplementary Figure 3). In order to correct for background (free CoASH and succ-CoA) in cell-free samples, as per the manufacturer’s instructions, CoASH Quencher and Quencher remover were used. Samples were then diluted with the reaction mix and fluorescence was measured (Excitation 535, Emission 589 nm) using a Clariostar plate reader (BMG, UK).

### 2.9 Nile Red plate assay

Nile Red plate assays for qualitative detection of PHAs were carried out as previously described (Kelwick et al., 2015b). Briefly, *E. coli* MG1655 transformed with either a negative control plasmid (EV_104) or a *phaCAB* operon (Native, C104, or C104 Δ(PhaC_C319A) were grown in 5 ml 2xYT+P media (supplemented with 34 μg/ml Chloramphenicol) overnight at 37°C with shaking (220 rpm). Liquid cultures were then streaked onto 2xYT+P-agar plates supplemented with 34 μg/ml Chloramphenicol, 0.5 μg/ml of Nile Red stain (Sigma-Aldrich, MO, USA, #72485-100MG) in 100% DMSO (v/v) and either 3% Glucose (w/v) or 120.48 g/L whey permeate. Nile Red plates were incubated for 24 h at 37°C and imaged using a Fuji Film LAS-5000 imager (Ex. 473 and Em. Cy5 filter).

### 2.10 Flow cytometry analysis of PHA production

Flow cytometry analysis of PHA content was carried out similarly to previous reports (Kelwick et al., 2015b; Lee et al., 2013). Briefly, *E. coli* MG1655 harbouring either a negative control plasmid (EV_104) or a *phaCAB* operon (Native, C104, or C104ΔPhaC_C319A) were grown overnight at 37°C in 5 ml of 2xYT+P medium supplemented with 34 μg/ml Chloramphenicol. The resultant cultures were diluted to an OD_600nm_ of 0.8 in 6 ml of PHA production media; 2xYT+P supplemented with 34 μg/ml Chloramphenicol and 120.48 g/L Molkolac instant demineralised whey permeate (Orchard Valley Food Ingredients, UK), and were cultured at 37°C for 24 hours. Subsequently, 1 ml of each culture was centrifuged (7,200 g), washed with 1 ml 1X PBS, and fixed with 35% ethanol [v/v] at room temperature for 15 minutes. Post-fixation, cultures were centrifuged (7,200 g), re-suspended in 1ml 1X PBS and stained with Nile Red (Sigma-Aldrich, MO, USA, #72485-100MG) to a final concentration of 20 μg/ml (in 100% DMSO) for 10 minutes on ice. Nile Red stained *E. coli* were diluted (1:100) into 1X PBS before being loaded into an Attune NxT (ThermoFisher Scientific, MA, USA) flow cytometer. PHA content was determined via flow cytometry analysis of Nile Red staining (YL2-A+, Ex. 560 nm, Em. 610 nm). Flow cytometry data analysis was carried out using FlowJo (v 10.1) software. Doublets were removed during gating. The background signal, as determined by the average geometric mean (YL2-A) of the appropriate, Nile Red stained, empty vector transformed *E. coli* was removed and these data were normalised to native-*phaCAB* engineered *E. coli*.

### 2.11 PHA purification

PHA purification was carried out using a scaled down version of a previously reported sodium hypochlorite-based method (Heinrich et al., 2012; Kelwick et al., 2015b). Briefly, glycerol stocks of *E. coli* MG1655 strains harbouring either a negative control plasmid (EV_104) or a phaCAB-operon (native, or C104) were used to inoculate flasks containing 30 ml of 2xYT+P medium (supplemented with 34 μg/ml Chloramphenicol). These starter cultures were grown overnight, with shaking (220 rpm), at 37°C. The resultant cultures were then diluted to an OD_600nm_ of 0.8 in 100 ml of PHA production media (2xYT+P media supplemented with 120.48 g/L whey permeate and 34 μg/ml Chloramphenicol (final concentration)). These PHA production cultures were then incubated at 37°C, with 220 rpm shaking, for 24 hours. Subsequently, 100 ml PHA production cultures were centrifuged at 4,200 rpm (Beckman J6-M1, USA) for 20 minutes. Post-centrifugation, bacterial cell pellets were re-suspended in 1X PBS, transferred into pre-weighed 50 ml tubes and centrifuged at 3,220 *g* for 15 minutes at 4°C. The supernatant from each 50 ml tube was removed and the cell pellets were dried at 70°C for 60 minutes and weighed. Dried cell pellets were re-suspended in 10 ml 1X PBS, centrifuged at 3,220 *g* for 15 minutes at 4°C, and the supernatant was discarded. Cell pellets were then suspended in 1X PBS with 1% Triton-X 100 (v/v in PBS) and then incubated for 30 minutes at room temperature. For the final PHA purification steps, cells were centrifuged at 3,220 *g* for 15 minutes at 4°C, washed with 1X PBS and then incubated with 12 ml aqueous sodium hypochlorite for 80 minutes at 30°C with 220 rpm shaking. The resultant purified PHA granules were centrifuged at 3,220 *g* for 30 minutes at room temperature, washed with distilled water and dried overnight at 37°C and then 2 hours at 70°C. To determine the weight of the dry cell pellets, the weight of the 50 ml tube was subtracted from the combined weight of the 50 ml tube and dry cell pellet. To determine the weight of the purified PHA, the weight of the 50 ml tube was subtracted from the combined weight of the 50 ml tube and PHA.

### 2.12 Monomer identification of *in vivo* produced PHAs using GC-MS

The P(3HB) and P(3HV) samples were subjected to methanolysis in a solution of 425 μl of methanol, 75 μl of sulphuric acid (6.6M) and 500 μl of dichloromethane in a small screw-top test tube and left to react for 3 hours at 100°C. After allowing the mixture to cool, 500 μL of dichloromethane and 1 ml of water were added; the mixture shaken vigorously for 1 minute followed by centrifugation at 4,000 rpm for 5 minutes. The organic phase was removed and transferred to a screw-cap glass vial. GC-MS analysis was performed on an Agilent Technologies 7890B GC and MSD 5977 series system with electron ionisation in scan mode. Helium was used as the carrier gas. The temperatures of the injector and MS transfer line were 240°C and 250°C respectively, whereas MS quadrupole and MS source were 150°C and 230°C respectively. The samples were analysed with an injection volume of 1 μl, at a split ratio of 14 to 1. A temperature program was used for efficient separation of the esters: 55°C for 3 minutes; temperature increase 15°C/minute until 200°C and hold for 3 minutes.

## 3. Results and discussion

### 3.1 Development of a whey permeate-based cell-free energy system

Conventional cell-free TX-TL reactions are broadly composed of three main components: a cell extract, an energy mix and the DNA construct (e.g. plasmid or linear PCR product) that encodes the protein, RNA/DNA based device or biosynthetic pathway that is being tested. Simplified, and/or automated cell extract preparation methods have led to improvements in cell-free performance (Krinsky et al., 2016; Kwon and Jewett, 2015; Sun et al., 2013). Likewise, changes in the composition and/or concentrations of cell-free energy mix components have also improved cell-free performance, largely through improved ATP regeneration, the recycling of inorganic phosphate and through the use of inexpensive single (e.g. glucose) or dual energy sources (e.g. maltose or maltodextrin and glutamate) that make optimal use of central metabolic pathways (e.g. glycolysis and tricarboxylic acid cycle) (Calhoun and Swartz, 2005; Caschera and Noireaux, 2015, 2014). Cell-free energy mixes can also be simplified through the rational testing and removal of unnecessary components that upon re-examination are largely historical artefacts from protocols that have now been superseded by improved cell-free methodologies. Such approaches have been used to develop minimal energy mixes that maintain cell-free performance but with seven fewer energy components than widely reported cell-free protocols (Cai et al., 2015).

Upon consideration of several cell-free methodologies, a fairly simplified cell extract preparation workflow was used in this study adapted from our previous cell-free work (Kelwick et al., 2016) and coupled with a recently reported glutamate-based minimal energy mix (Cai et al., 2015). Using these cell-free methods several batches of *E. coli* MG1655 TX-TL cell extracts and minimal energy mix were generated (see materials and methods 2.3 and 2.4). These cell-free reactions were customised for the purposes of characterising PHA biosynthetic pathways through the inclusion of whey permeate. Whey permeate is a waste by-product of industrial cheese production that has been proposed as an attractive low-cost feedstock for microbial PHAs production *in vivo* - particularly P(3HB) (Bosco and Chiampo, 2010; Ryan and Walsh, 2016). Whey permeate contains a high percentage of lactose (>70% of the mass), which can be readily metabolised *in vivo* by *E. coli* MG1655, using its β-galactosidase enzyme, to generate glucose for glycolytic processing into Acetyl-CoA (Pasotti et al., 2017). Acetyl-CoA can then directly feed into *phaCAB* biosynthetic operons where it is enzymatically processed by PhaA (3-ketothiolase) to form acetoacetyl-CoA. Subsequently, PhaB (acetoacetyl-CoA reductase) reduces acetoacetyl-CoA to form (R)-3-hydroxybutyl-CoA ((R)-3HB-CoA), which is finally polymerised by PhaC (PHA synthase) to form the final PHA polymer - poly-3-hydroxybutyrate (P(3HB)).

To determine whether whey permeate could be similarly metabolised *in vitro* a series of *E. coli* MG1655 cell-free TX-TL reactions were setup that included a minimal energy mix but with different concentrations of whey permeate (0-5.748 g/L) and a constitutive *gfpmut3b* expression plasmid (101_GFP) (Figure 1a). Cell-free TX-TL GFPmut3b production data are representative of three different cell extract batches (Figure 1). An endpoint (10 h) analysis of cell-free GFPmut3b production revealed that whey concentrations of at least 0.064 g/L could either partially or completely inhibit cell-free TX-TL activity (Figure 1b). It is likely that relatively high concentrations of whey permeate salts are responsible for this inhibitory effect (Supplementary Table 3). Indeed, previous reports have shown that the concentrations of different salts can significantly impact cell-free performance (Ge et al., 2011). We therefore used demineralised whey permeate which has been processed to lower its salt and mineral content. Lower concentrations of whey permeate were not inhibitory and interestingly 0.004 g/L of whey permeate enhanced relative cell-free GFPmut3b production by ~20% ± 7.5 (Figure 1b). Cell-free dynamics, in terms of the rate of cell-free GFPmut3b production, were also enhanced in cell-free reactions with 0.004 g/L of whey permeate in comparison to reactions that included the standard minimal energy mix (0 g/L) (Figure 1c).

**Figure 1.**
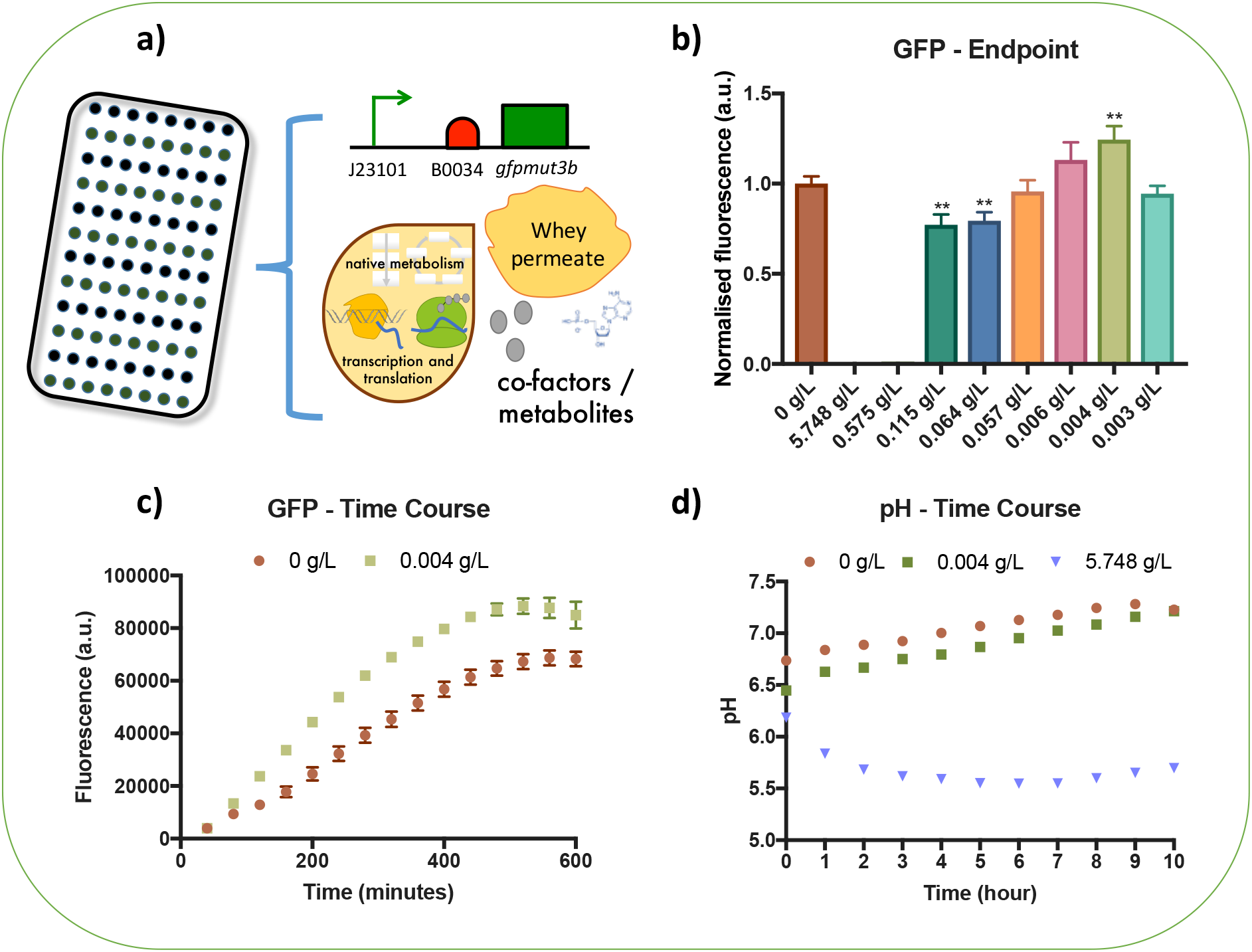
Whey permeate enhances cell-free transcription-translation activity likely through glycolysis. **(a)** Schematic depicts whey permeate containing cell-free transcription-translation reactions that were used to test *in vitro* GFP protein production, **(b)** Endpoints (10 h) of cell-free reactions containing GFP expression plasmid (101_GFP) and 0 to 5.748 g/L whey permeate as indicated. The background fluorescence of cell-free reactions using negative control plasmid (EV101) were subtracted and these data were normalised to the relative fluorescence of cell-free reactions containing 0 g/L whey permeate, **(c)** Time-course analysis of GFP production in cell-free reactions which included GFP expression plasmid (101_GFP) and 0 or 0.004 g/L of whey permeate. The background fluorescence of cell-free reactions using negative control plasmid (EV101) were subtracted, **(d)** Time-course analysis of pH in cell-free reactions which included the negative control plasmid (EV101) and 0–5.748 g/L of whey permeate. Error bars denote standard error of the mean. Student t-test, **P<0.01.

The metabolism of whey permeate through glycolysis, is likely to lead to an increase in the production of lactate and acetate which have been previously shown to decrease the pH of cell-free TX-TL reactions (Kara and James, 2005). Previous reports have also shown that the coupling of maltose with glycolytic processing not only decreases the pH of cell-free reactions, through the production of lactate and acetate, but beneficially, enhanced glycolytic activity also increases ATP generation (Caschera and Noireaux, 2014). The pH of cell-free reactions can be measured using a pH sensitive fluorescent dye - known as SNARF-5F (see materials and methods 2.6). Similar to a previous report, we setup a SNARF-5F calibration curve to convert between dye fluorescence and pH (Supplementary Figure 1) (Caschera and Noireaux, 2014). The pH of cell-free reactions without whey permeate (0 g/L) was generally stable throughout the 10 h reaction (Figure 1d). Whereas the pH was slightly lower in cell-free TX-TL reactions with 0.004 g/L of whey permeate and the highest concentration tested (5.748 g/L) resulted in a significant pH drop from 6.2 to 5.6 (Figure 1d). Thus, an optimal whey permeate concentration is 0.004 g/L, which we found enhances *in vitro* protein production likely through increased ATP generation as a consequence glycolytic metabolism of lactose/glucose. Whey permeate was therefore included in subsequent prototyping experiments.

### 3.2 Cell-free TX-TL characterisation of PHAs biosynthetic operons

In our previous study, the native *phaCAB* operon from *Cupriavidus necator* was used to engineer several *phaCAB* operon variants that enhanced *in vivo* PHAs production in *E. coli* (Kelwick et al., 2015b). Amongst these variants, the C104 constitutive *phaCAB* operon (BBa_K1149052) was designed such that the native promoter and RBS were replaced by an Anderson constitutive promoter (BBa_J23104) and a strong synthetic ribosome binding site (RBS - BBa_B0034) (supplementary file 1). Previously, the C104 design enhanced *in vivo* P(3HB) production from glucose by up to three-fold in comparison to native *phaCAB*-engineered *E. coli* (Kelwick et al., 2015b). Since, the native and C104 *phaCAB* operons produce different levels of P(3HB) *in vivo*, several cell-free TX-TL experiments were designed to determine whether the differences in operon activity could be determined *in vitro* (cell-free).

Initial attempts to detect the cell-free TX-TL production of P(3HB) were based on several methods adapted from previously described liquid chromatography–mass spectrometry (LC-MS) and GC-MS approaches (de Rijk et al., 2005). However, these methods were found to be unreliable for detecting P(3HB) in cell-free TX-TL reactions. There are several reasons which could explain a lack of detection. Firstly, established PHA analyses methods were largely designed for detecting highly concentrated and purified, high molecular weight PHAs polymers (Huang and Reusch, 1996). Whereas, small-scale cell-free TX-TL prototyping reactions likely generate relatively low concentrations of PHAs at a range of molecular weights (polymer chain lengths). Secondly, the samples also contain a complex background of many different cell extract-derived metabolites that further complicate PHAs detection. Furthermore, additional processing steps that are required to depolymerise P(3HB) into the more easily detectable monomer form (3HB) may increase the risk of sample loss. Due to these challenges, additional methods development was carried out.

*E. coli* MG1655 cell extracts were spiked with known concentrations of commercially available 3-hydroxybutyrate (3HB) and these spiked extracts were analysed using different GC-MS methodologies. From these experiments, an optimised GC-MS method was developed, along with a calibration curve, that was used to detect 3HB in the low micromolar range within TX-TL cell extracts (Materials and Methods 2.7; Supplementary Figure 2). Method optimisation was achieved through improvements in the temperature gradient for better separation of derivatised 3HB from the cell extract background. A more sensitive and selective mass spectrometry (MS) acquisition was also acquired, using the selected ion monitoring (SIM) approach, coupled with monitoring m/z values, as described in the methods section 2.7, during the GC-MS analysis. In parallel we also engineered an additional *phaCAB* operon which maintained the same operon structure as C104, but with an inactive PhaC enzyme which is unable to polymerise (R)-3HB-CoA into P(3HB). Site specific mutation of the PhaC catalytic Cys 319 with alanine gave a C104 Δ(PhaC_C319A) *phaCAB* operon (materials and methods 2.2). This additional operon reduced the sample processing steps needed for PHA polymers and allowed the direct detection of PHAs monomers produced in cell-free reactions.

Cell-free TX-TL reactions were setup and analysed using our optimised GC-MS method and these data are representative of three cell extract batches. GC-MS analysis confirmed an average 3HB production of 3.15 μM ±1.5 in C104Δ(PhaC_C319A) cell-free TX-TL reactions and 3HB was not detectable in negative control reactions (EV104) (Figure 1b).

Cell-free TX-TL reactions can also be coupled with other types of assays that can provide additional insights into the cell-free activities of PHA biosynthetic operons. For instance, a pH analysis of negative control (EV104) and C104 *phaCAB* operon TX-TL reactions (Figure 2c) confirmed that, as expected, the pH of these cell-free reactions is largely determined by whey permeate metabolism and not by the activities of the PhaCAB enzymes. We also hypothesised that Acetyl-CoA content would be lower in TX-TL reactions that included a plasmid encoding a *phaCAB* operon, relative to the negative control (EV104). We reasoned that production of PhaCAB enzymes would more rapidly reduce Acetyl-CoA pools through conversion towards P(3HB). To test this, we set up a series of *E. coli* MG1655 TX-TL cell-free reactions and assayed Acetyl-CoA endpoint content as described in materials and methods. The average endpoint (1 h) Acetyl-CoA content was higher in negative control reactions (EV104 8.17 pmol/μl ±3.78) than reactions with either a native (3.19 pmol/μl ±1.42), C104 (0.55 pmol/μl ±0.55) or C104Δ(PhaC_C319A) (0.57 pmol/μl ±0.57) *phaCAB* operon. Together, these data demonstrate that PHAs can be produced and detected within cell-free TX-TL reactions and despite some variation between Acetyl-CoA assay, these types of cell-free analyses could be used to screen for differences in the activities of different *phaCAB* operons. Indeed, the general pattern in the Acetyl-CoA assays indicate that the C104 *phaCAB* operon is likely to enhance *in vivo* PHAs production, relative to the native *phaCAB* operon (Kelwick et al., 2015b)(Figure 4). Likewise, these data also demonstrate that coupling cell-free TX-TL systems with an industrial feedstock (whey permeate in our case) can potentially enhance *in vitro* protein production (Figure 1) and enable the prototyping of PHA biosynthetic operons that relate to the metabolic context of *in vivo* PHA production.

**Figure 2.**
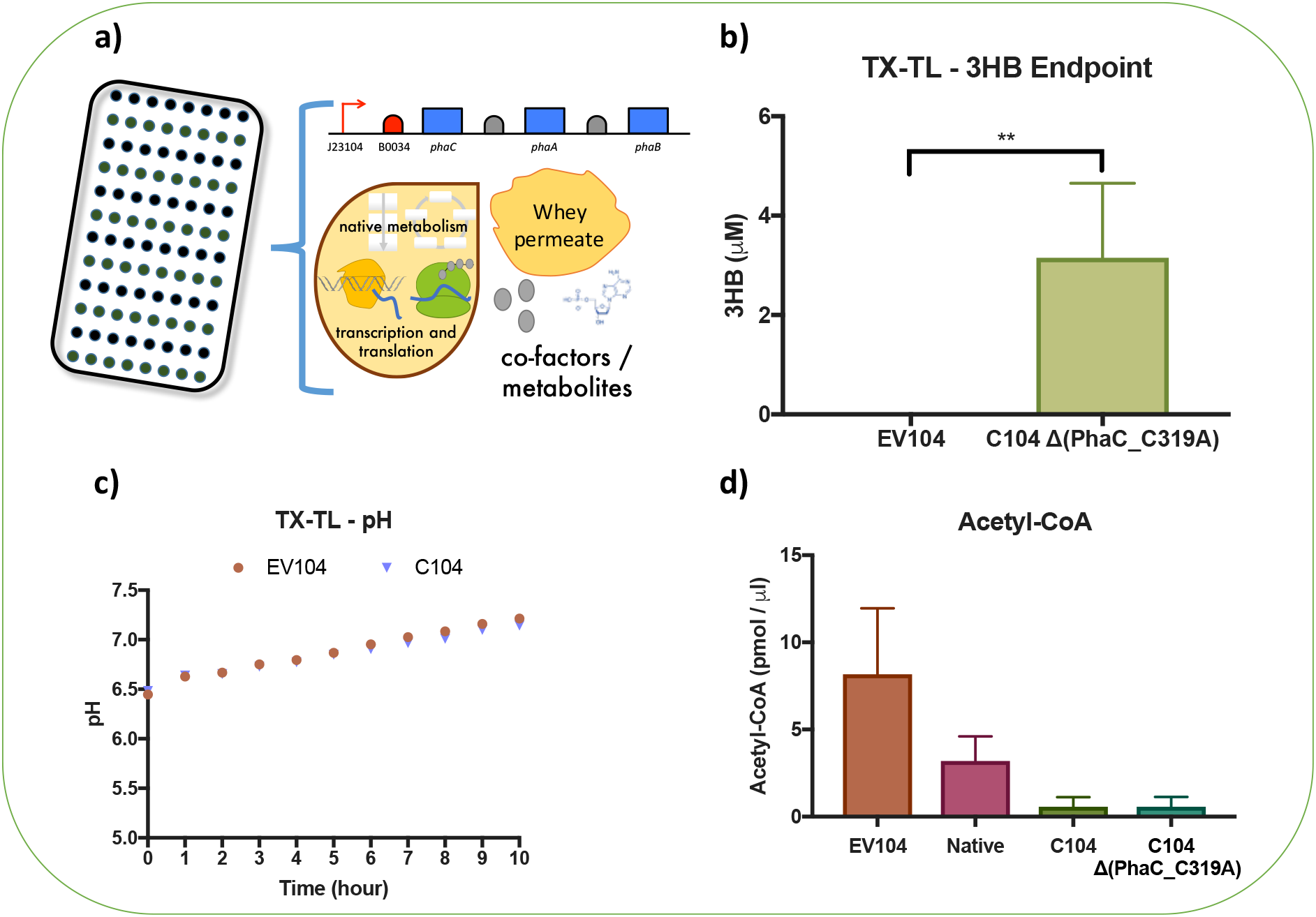
Cell-free TX-TL characterisation of *phaCAB* biosynthetic operons and PHAs production from whey permeate. **(a)** Schematic depicts the whey permeate containing cell-free transcription-translation (TX-TL) reactions used to test the cell-free production of 3-hydroxybutyrate (3HB). **(b)** Endpoint (5 h) GC-MS analysis of 3HB production in whey permeate containing cell-free reactions. These reactions included either a negative control plasmid (EV104) or a plasmid encoding a *phaCAB* operon with an inactive PhaC (C104 Δ(PhaC_C319A)). **(c)** Time-course analysis of pH in whey permeate containing cell-free reactions that included either a negative control plasmid (EV104) or a plasmid encoding a *phaCAB* operon (C104). **(d)** Endpoint (1 h) analysis of Acetyl-CoA content in whey permeate containing cell-free reactions. These cell-free reactions included either a negative control plasmid (EV104) or a plasmid encoding a *phaCAB* operon (Native, C104 or C104 Δ(PhaC_C319A)). Error bars denote standard error of the mean. Student t-test, **P<0.01.

### 3.2 Biotransformation of whey permeate in *phaCAB*-engineered cell extracts

Extract-based cell-free metabolic engineering (also known as cell-free protein synthesis driven metabolic engineering – CFPS-ME) can be categorised into two main strategies; one that makes use of cell-free transcription and translation systems and the other involving mixing cell extracts (Karim and Jewett, 2016). Such strategies use cell-free TX-TL systems for *in vitro* enzyme production and expression of entire biosynthetic pathways. Cell-free metabolic engineering can also be used to construct biosynthetic pathways through the rational mixing of different cell extracts (lysates) that include the necessary enzymes, co-factors and metabolic pathways. It is possible to combine these different cell-free strategies. For instance, bacterial strains that produce desirable enzymes during cell growth can later be lysed and processed to generate cell extracts for use in cell-free transcription-translation reactions. Therefore, a combined cell-free metabolic engineering approach could be used as a prototyping platform for the *in vitro* characterisation of additional enzymes that may directly or indirectly enhance PHAs production (e.g. metabolite recycling enzymes).

In order to test the concept of cell-free biotransformation reactions, several batches of cell extract from C104 Δ(PhaC_C319A) *phaCAB*-engineered *E. coli* MG1655 (S_RK004) were generated. These cell extracts include endogenous β-galactosidase, central metabolic pathways (e.g. glycolysis) and all three PhaCAB enzymes (inactive PhaC), and thus contain all the enzymes needed for the biotransformation of whey permeate into the PHA monomers - (R)-3HB-CoA (Figure 3a). Biotransformation reactions were carried out as described in materials and methods. Endpoint (5 h) GC-MS analyses of the cell-free biotransformation reactions revealed that average (R)-3HB-CoA content was 8.84 μM ±2.47 in reactions with 0 g/L of whey permeate and 32.87 μM ±6.58 in reactions with 0.004 g/L of whey permeate (Figure 3b). Even those cell-free biotransformation reactions without an exogenously added glycolysis substrate (0 g/L whey permeate), produced (R)-3HB-CoA (Figure 3b), suggesting that these cell extracts generated Acetyl-CoA using an alternative pathway, such as through β-oxidation (Tee et al., 2014). Interestingly, cell-free production of (*R*)-3HB-CoA from whey permeate (0.004 g/L) was around 10x higher in biotransformation reactions (Figure 3b) than in TX-TL reactions (Figure 2a). We anticipate that, with further developments in strain engineering, cell-free biotransformation reactions could conceivably be used for PHAs production. Of course, cell-free biotransformation reactions are not limited to PHAs production applications and can potentially be combined with cell-free TX-TL approaches for screening enzymes that might enhance PHAs production in both cell-free and *in vivo* contexts.

**Figure 3.**
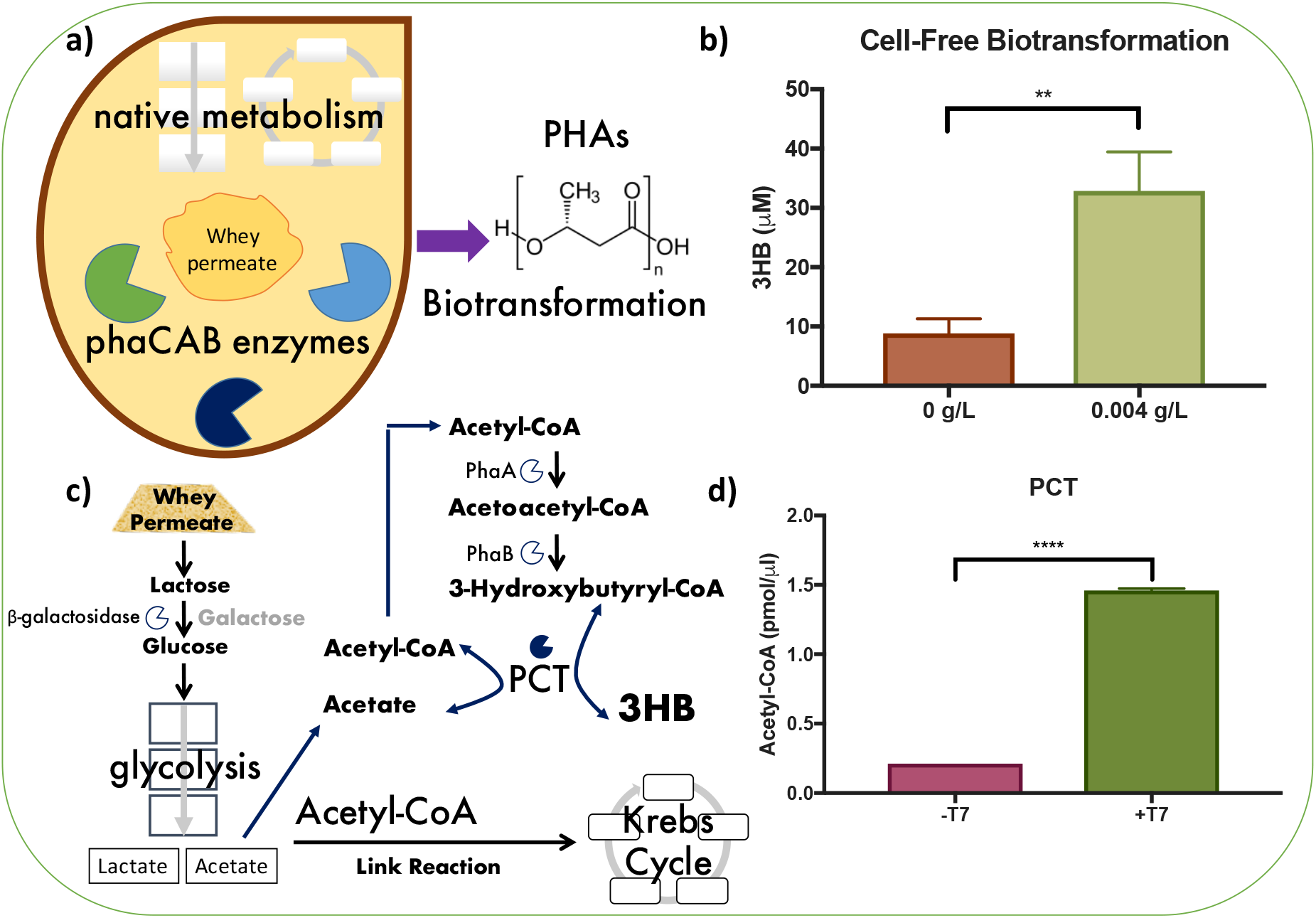
Biotransformation of whey permeate into 3-hydroxybutyrate (3HB) in *phaCAB*-engineered *E. coli* cell extracts. **(a)** Schematic depicts the cell-free biotransformation of whey permeate into 3-hydroxybutyrate (3HB) using engineered cell extracts that contain PhaCAB enzymes. In these extracts, the PhaC enzyme was engineered to be inactive (PhaCΔC319A) to ensure that cell-free produced 3HB was not polymerised, **(b)** Endpoint (5 h) GC-MS analysis of 3HB production in cell-free biotransformation reactions. These cell-free reactions included either 0 or 0.004 g/L of whey permeate, **(c)** Schematic depicts the Acetyl-CoA recycling activity of propionyl CoA transferase (PCT) within the context of whey permeate metabolism in cell extracts. (d) End point (2.5 h) analysis of Acetyl-CoA content in whey permeate containing cell-free biotransformation reactions that also included plasmids encoding T7 inducible expression of an Acetyl-CoA recycling enzyme – PCT. Thus, extracts with T7 RNA polymerase (+T7) also enable transcription and translation of the PCT enzyme. Error bars denote standard error of the mean. Student t-test, **P<0.01 and ****P<0.0001.

As an exemplar, we tested the cell-free expression of the propionyl CoA transferase (*pct*) gene from *Clostridium propionicum* in C104 Δ(PhaC_C319A) *phaCAB*-engineered cell extracts. PCT couples the release of CoA from (R)-3HB-CoA with the generation of Acetyl-CoA from acetate (Figure 3c) (Ken’ichiro Matsumoto et al., 2013a; Selmer et al., 2002). Since, whey permeate is likely to increase the glycolytic production of acetate and PCT reactions generate Acetyl-CoA (a precursor of (*R*)-3HB-CoA), it was anticipated that Acetyl-CoA and 3HB levels would be relatively higher in cell-free biotransformation reactions that express the *pct* gene. The *C. propionicum pct* gene sequence was sourced from Genbank (AJ276553.1) and cloned into a T7 expression plasmid (as described in materials and methods 2.2) which was termed T7_PCT (pRK12). Combined biotransformation/TX-TL cell-free reactions were setup with C104 Δ(PhaC_C319A) *phaCAB*-engineered cell extracts, in combination with the T7_PCT plasmid, and either 0 (-T7) or 25 (+T7) units of T7 RNA polymerase (materials and methods 2.5). Thus, only those cell-free reactions that contain T7 RNA polymerase will express the *pct* gene. Endpoint analyses revealed that average Acetyl-CoA content was higher in cell-free reactions that contained T7 RNA polymerase (1.46 pmol/μl ±0.01) in comparison to control reactions with no T7 RNA polymerase (0.21 pmol/μl ±0.00) (Figure 3d). Whilst the PCT driven increase in Acetyl-CoA is relatively modest and did not result in increased (R)-3HB-CoA content (data not shown), these data suggest that coupled cell-free biotransformation and TX-TL prototyping could be useful in helping to identify metabolite recycling enzymes that might improve PHAs production *in vivo*. Indeed, in other experimental contexts *pct* genes from both *C. propionicum* and *Megasphaera elsdenii* have been shown to enhance *in vivo* Acetyl-CoA content (Ken’ichiro Matsumoto et al., 2013a, 2013b; Selmer et al., 2002; Tajima et al., 2016).

### 3.3 *In vivo* characterisation of PHAs production from whey permeate

Finally, *in vivo* PHA production experiments were carried out using the *phaCAB*-engineered *E. coli* MG1655 strains and negative control. The optimal whey concentration for cell-free assays (0.004 g/L, Figure 1b) is likely to be insufficient for *in vivo* PHAs production, where a relative excess of carbon is desirable. Therefore, a previously described whey permeate concentration of 120.48 g/L, corresponding to 100 g/L of lactose, was used in all *in vivo* assays (Bosco and Chiampo, 2010).

Qualitative analysis of Nile Red stained E. *coli* confirmed PHAs polymer production in native, and C104 *phaCAB*-engineered *E. coli* (Materials and methods 2.9, Figure 4 and Supplementary Figure 4). Whereas, both negative control (EV104) and C104 Δ(PhaC_C319A) *phaCAB*-engineered *E. coli* did not generate PHAs granules (Figure 4 and Supplementary Figure 4). In addition to the plate assays, a semi-quantitative Nile Red flow cytometry analysis of *in vivo* PHAs production was also carried out (Materials and methods 2.10, Figure 4b, Supplementary Figure 5, Supplementary Figure 6 and Supplementary Table 4). For these analyses, a gating strategy was implemented that excluded cell doublets and distinguished between negatively and positively Nile Red stained cell populations (Supplementary Figure 5 and Supplementary Table 4). Post gating, the background signal of negative control cells (EV104) was removed and these data were normalised to the Nile Red fluorescence of native *phaCAB*-engineered *E. coli*. Nile Red fluorescence levels were on average 2.18 ±0.05 fold-higher in C104 *phaCAB*-engineered *E. coli* than native *phaCAB*-engineered *E. coli* levels (Figure 4b and Supplementary Figure 6) indicating that PHAs content is higher in C104 *phaCAB*-engineered *E. coli*.

**Figure 4.**
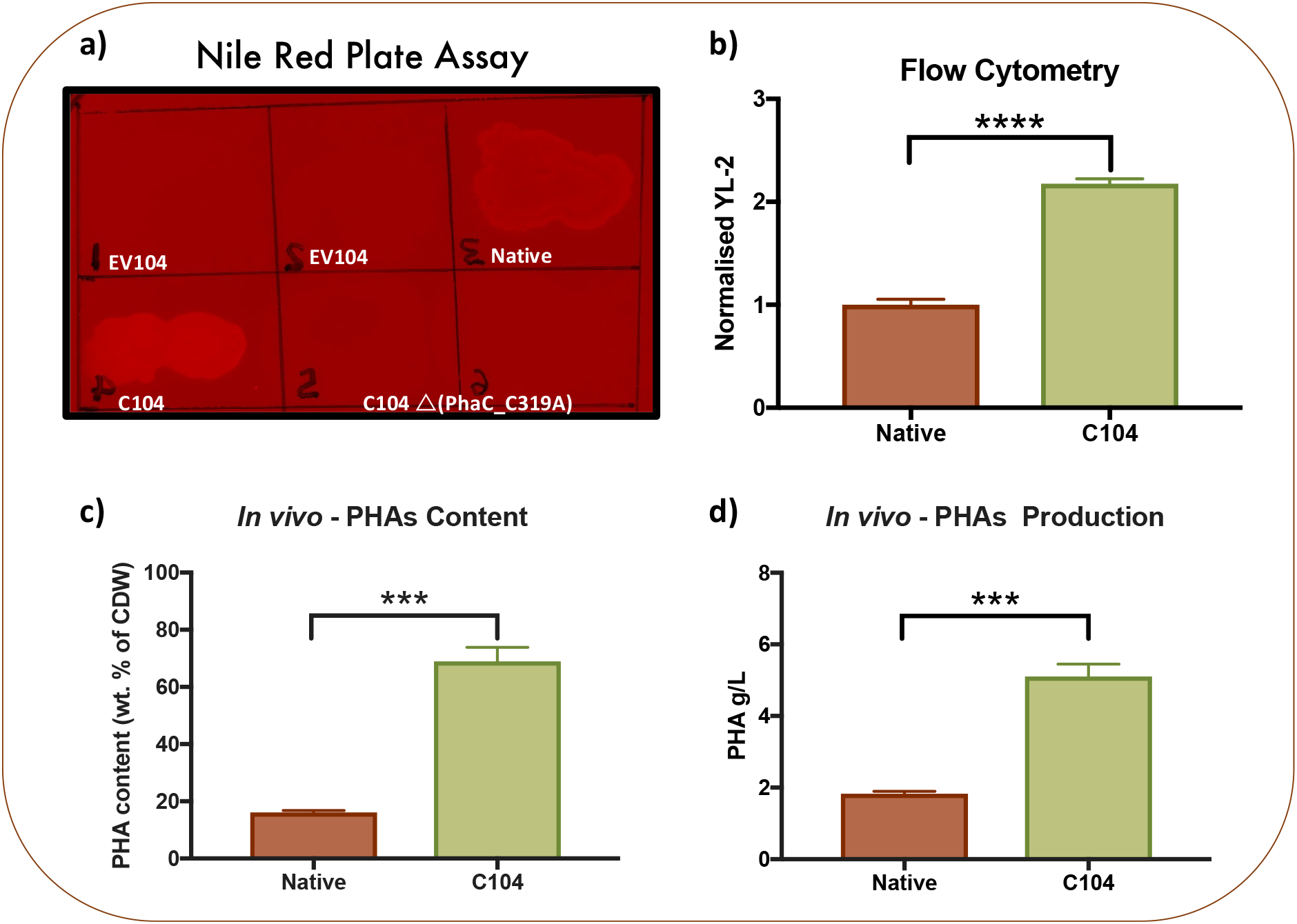
*In vivo* Characterisation of PHAs production from whey permeate in *phaCAB*-engineered *E. coli*. **(a)** *E. coli* MG1655 transformed with either a negative control plasmid (EV_104) or a *phaCAB* operon (native, C104, or C104ΔPhaC_C319A) were streaked onto 2xYT+P-agar plates supplemented with 3% glucose (w/v), 34 μg/ml Chloramphenicol and 0.5 μg/ml (final concentration) of Nile Red stain. Nile Red plates were incubated for 24 h at 37°C and then imaged using a Fuji Film LAS-5000 imager (Ex. 473 and Em. Cy5 filter), **(b)** *E. coli* MG1655 transformed with either an empty vector (EV104) or a *phaCAB* operon (native or C104) were Nile Red stained and analysed via flow cytometry (Attune NxT - YL2-A, Ex. 560 nm, Em. 610 nm). The background signal, as determined by the average geometric mean (YL2-A) of Nile Red stained, empty vector (EV_104) transformed *E. coli* was removed and these data were normalised to native *phaCAB*-engineered *E. coli*. In order to determine PHA production in *phaCAB*-engineered *E. coli* (native and C104), 100 ml PHA production cultures (2xYT+P media - supplemented with 120.48 g/L milk whey permeate and 34 μg/ml Chloramphenicol) were setup, as described in materials and methods, and were cultured for 24 hours at 37°C with shaking (220 rpm). PHAs were purified from these PHA production cultures and measured as **(c)** PHA content (weight [wt] *%* of cell dry weight [CDW]) and **(d)** PHAs production (g/L). Error bars indicate the standard error of the mean. Student t-test, ***P<0.001, and ****P<0.0001.

*In vivo* PHAs production experiments involving *phaCAB*-engineered *E. coli* were carried out in whey permeate production media and post-production, a sodium hypochlorite-based method was used to purify PHA content from these strains (materials and methods 2.11). PHAs production from whey permeate was almost three times higher in C104 *phaCAB*-engineered (5.10 ±0.35 g/L) cultures than native PHAs production cultures (1.83 ±0.07 g/L) (Figure 4c and Supplementary Table 5). PHAs production was also validated using GC-MS (Materials and methods 2.12, Supplementary Table 6). Similarly, PHAs content expressed as % weight of cell dry mass was also significantly higher in C104 *phaCAB*-engineered cultures (68.96 % ±4.91) than native PHAs production cultures (16.14 % ±0.68) (Figure 4d and Supplementary Table 5). Although the reaction scales, whey concentrations and PHAs production yields differ between these *in vivo* and *in vitro* cell-free assays, it is clear that the C104 *phaCAB* operon, and its derivative C104 Δ(PhaC_C319A) were more active than the native operon as observed in both cell-free and *in vivo* assays. This therefore illustrates how cell-free TX-TL approaches can complement *in vivo* workflows for prototyping PHA biosynthetic operons.

## 4. Summary and Conclusions

Despite the environmental consequences that are associated with the mass production of oil-derived plastics, global demand is likely to continue to increase unless viable economic alternatives are developed (Eriksen et al., 2014; Geyer et al., 2017). The polyhydroxyalkanoates are a family of biodegradable biopolymers, that could potentially be used as sustainable alternatives to replace several widely used oil-derived plastics (e.g. polypropylene). However, PHAs are currently more expensive to produce than oil-derived plastics which has hampered their adoption and therefore, more efficient PHAs production processes would be desirable. An array of advancements in synthetic biology and metabolic engineering have already led to improvements in PHAs production though we anticipate that cell-free metabolic engineering approaches, several of which have been underutilised in PHAs research (e.g. cell-free TX-TL), could lead to additional innovations in PHAs production strategies. Indeed, cell-free metabolic engineering approaches have already been used to prototype biosynthetic pathways in other contexts (Karim and Jewett, 2016). Yet, interestingly, cell-free TX-TL systems have not previously been used to prototype PHA biosynthetic operons. This may in part relate to the challenges associated with detecting PHAs and PHA monomers produced in small scale cell-free reactions. In order to overcome this and to accelerate the adoption of cell-free TX-TL prototyping platforms in PHAs research, several cell-free TX-TL approaches for characterising PHA biosynthetic pathways were developed. *E. coli* MG1655 cell-free prototyping reactions were customised with whey permeate as an energy source, an industrial waste that has been previously used as a low-cost feedstock for optimising *in vivo* PHAs production. The inclusion of an optimal concentration of whey permeate enhanced relative *in vitro* protein production and subsequent experiments also demonstrated, for the first time, the production and GC-MS detection of 3-HB (a PHAs monomer) within cell-free TX-TL reactions. pH and Acetyl-CoA assays were also used to demonstrate that additional insights into the activities of PHA biosynthetic pathways can also be gained through carrying out cell-free TX-TL prototyping assays. Therefore, these data demonstrate that suitably customised cell-free TX-TL systems can be used to characterise PHA biosynthetic operons within a metabolic context that relates to *in vivo* production. Additionally, we also demonstrated that coupled cell-free biotransformation/TX-TL strategies can be used to screen for useful metabolite recycling enzymes for enhancing *in vivo* PHAs production. As an exemplar, we *in vitro* expressed and characterised an Acetyl-CoA recycling enzyme (*pct*) within *phaCAB*-engineered biotransformation cell extracts.

More broadly we envision that these types of cell-free metabolic engineering approaches could conceivably be used in combination with *in vivo* strategies for optimising PHAs production (Figure 5). Ultimately, the continuing development of cell-free TX-TL metabolic engineering approaches may lead to desirable innovations in PHAs production that enhance their potential as sustainable alternatives to oil-derived plastics.

**Figure 5.**
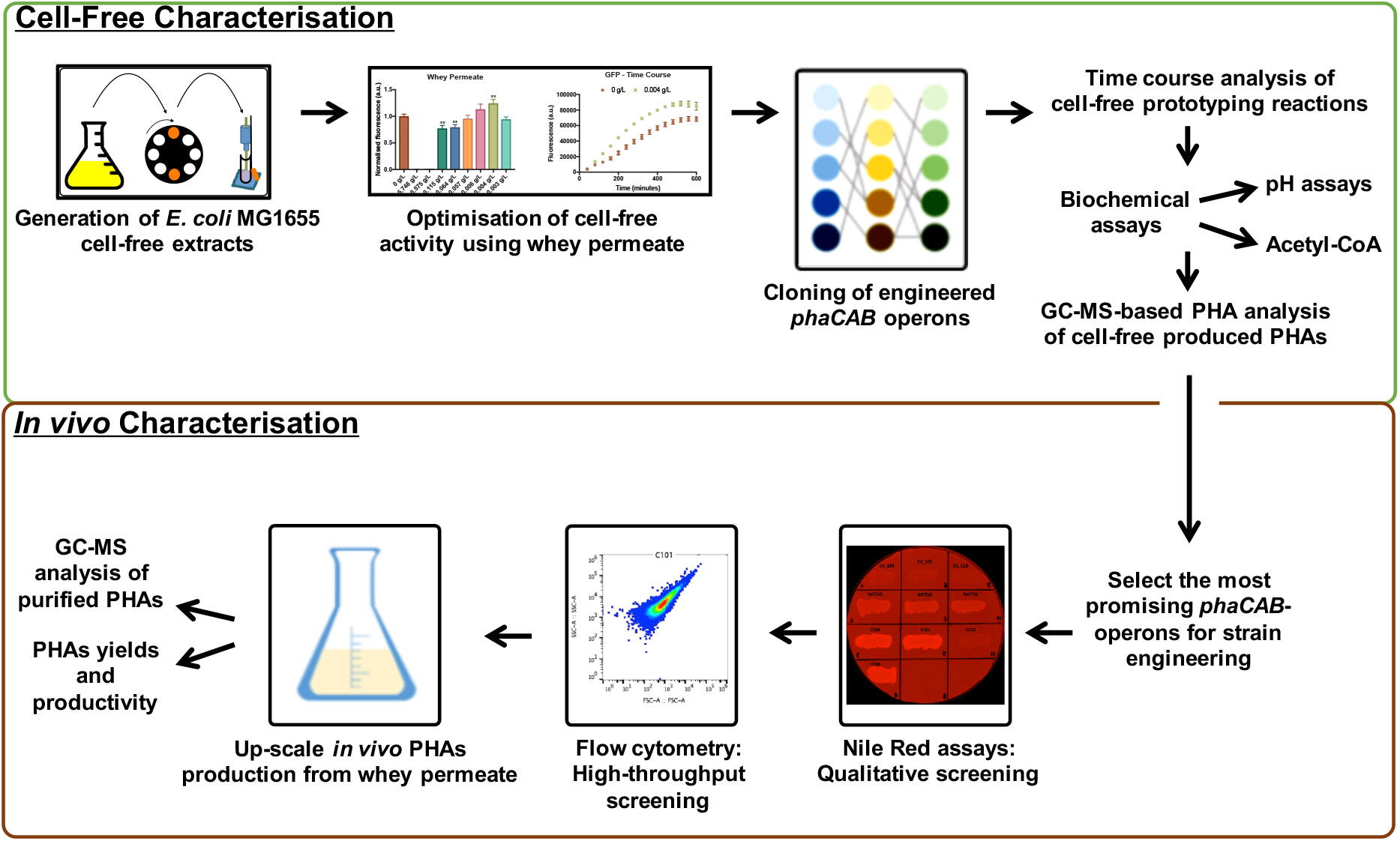
Cell-free and *in vivo* characterisation of PHAs production. Cell-free prototyping approaches complement *in vivo* strategies for characterising Polyhydroxyalkanoates (PHAs) production from whey permeate.

## Author Contributions

Project conception (R.K. and P.S.F.), designed the experiments (R.K. and L.R.), performed the experiments (R.K. and L.R.) and analysed the data (R.K. and L.R.). GC-MS and LC-MS assays and analysis (S.M.C. D.B. and R.K.). Wrote the manuscript (R.K.). Edited and reviewed the manuscript (R.K., L.R. A.W. P.S.F. S.M.C. and D.B.). Project support and advice (P.S.F. and A. W.).

## Acknowledgements

We wish to acknowledge the support of the Engineering and Physical Science Research Council (EPSRC) – [EP/J02175X/1] and that of our colleagues in the Centre for Synthetic Biology and Innovation (CSynBI), SynbiCITE at Imperial College London and also colleagues from The Flowers Consortium. We would also thank the Erasmus+ programme for supporting Luca during his master’s degree placement project.

## Data availability

Data available in supplementary material.

## Supplementary data

Supplementary File 1

